# Physicochemical and microbiome changes in queso crema de Chiapas during ripening

**DOI:** 10.1101/2025.04.02.646936

**Authors:** Blanca Nayelli Ocampo Morales, Arturo Hernández Montes, Karel Estrada, Ernestina Valadez Moctezuma

**Author notes:** (KE); (EVM).

## Abstract

The dynamic transformations of physicochemical, microbiological, and metagenomic profiles in Queso Crema de Chiapas were studied during distinct ripening stages (2, 29, and 58 days). While the fundamental physicochemical characteristics—including protein, fat, and salt content—remained remarkably stable, a distinctive evolution in microbial diversity was observed, characterized by a decline in bacterial genera and a concurrent increase in fungi and yeasts as maturation progressed.

The most abundant bacterial genera were *Streptococcus*, *Lactobacillus*, and *Lactococcus* throughout ripening. *Streptococcus* and *Lactobacillus* increased as ripening time progressed, while Lactococcus exhibited an opposite trend. In addition, several predominant fungal species were identified across the three ripening periods, including *Candida versatilis*, *Candida etchellsii*, and *Candida tropicalis*.

*Candida etchelsii* decreased, and *Candida tropicalis* increased with ripening time. Notably, low levels of potentially pathogenic microorganisms were detected. This study highlights the influence of ripening duration on microbial composition, providing valuable insights for the production of artisanal cheeses and the enhancement of their quality.

## Introducción

Several artisanal cheeses are produced in Mexico, including “Queso Crema de Chiapas”. The production of this cheese is differentiated by the resting of raw milk, prolonged fermentation of the paste, and kneading (1). As ripening occurs, the cheese can have a soft, semi-hard, or hard texture (2). For sale, it is presented in rectangular shape wrapped in plastic, aluminum foil, and cellophane. The cheese can be presented as a natural paste or occasionally seasoned paste and is consumed from one week of processing to more than three months of ripening (3). On the other hand, it has been explored that this cheese is a source of bioactive compounds such as: antioxidants, angiotensin-converting enzyme (ACE) inhibitors, and antimicrobial activity (4); (5); (6). Lactic acid bacteria (LAB) and fungi are responsible for desirable sensory characteristics such as texture, aromas, and flavors (7). However, the hydrolysis of peptides by bacteria gives rise to sour, sweet, or even unpleasant flavors. By breaking down proteins and fats, fungi also generate various volatile compounds (8). The microorganisms present in cheese can be modified because ripening and salt concentration affect their physicochemical properties (9). Milking conditions, herd feeding, manufacturing process, storage conditions, and interactions between cheese components-microorganisms and microorganisms-microorganisms also play a role in microbial dynamics (10). In the present study, Queso Crema de Chiapas from Mexico was characterized by physicochemical and microbiological tests and molecular identification of microorganisms during ripening.

## Materials and methods

### Obtaining samples of Queso Crema de Chiapas

The cheeses were made from raw bovine milk and three pieces were collected from each of the three batches made on consecutive days and left to mature for different times: T1 = 2, T2 = 29 and T3 = 58 days (d), with three replicates (n = 3) per condition. Samples were obtained from a cheese factory located at 15° 43’ 39.9354“ north latitude and -93° 15’ 45.4314” west latitude in the municipality of Pijijiapan, Chiapas, Mexico. The samples were transported at room temperature to the laboratory.

### Physicochemical analysis of Queso Crema de Chiapas

The physicochemical analysis considered the determination of protein, fat, total solids, and NaCl using the FoodScanTM Lab equipment (FOSS Analytical AB, Hillerød, Denmark); moisture (11) and ash (12) were quantified by gravimetric method. Each of the measurements was performed in triplicate. On the other hand, the Aqualab Series 3 instrument (Decagon Devices Inc., Washington, USA) was used for a_w_ determination (13).

Carbohydrate (CHO’S) estimation was obtained by difference [CHO’S (%) = Total Solids (TS) %-(fat %+protein %+ash %)](14).

### Microbiological analysis

For microbiological analyses, six blocks of cheese (n=6) from three different lots were used, and two replicates of each; a 10 g portion was taken from the inside of each block (15) and homogenized with 90 mL of sterile 0.1 % peptone wáter (16). Aerobic mesophilic bacteria counts were performed according to NOM-092-SSA1-1994. For fungi and yeasts, NOM-111-SSA1-1994 was followed and total coliform bacteria were quantified using NOM-113-SSA1-1994.

### Identification of bacteria, fungi, and yeasts by metagenomics

The V3 and V4 regions of the 16S gene were sequenced to identify bacteria, while the internal transcribed spacers ITS1-ITS2 were sequenced to identify fungi and yeasts. Sequencing was performed on the Illumina MiSeq platform.

### DNA extraction

DNA was extracted from cheeses at three ripening times with two replicates. Replicates belonging to the same ripening time were mixed. Samples were ground in liquid nitrogen until very fine particles were obtained. Then, 0.1 g of cheese was weighed and the extraction protocol of the ZymoBIOMICS DNA kit (California, USA) was followed. Subsequently, it was quantified with a UVS-99 nanoDrop (Avans Biotechnology, Taipei, Taiwan, China) at 260 nm to determine DNA purity and integrity.

## Statistical analysis

### Physicochemical and microbiological statistical analysis

The Kolmogorov-Smirnov normality test was applied using the NCSS 2007 program (NCSS LLC, Kaysville, Utah, USA) for the variables a_w_, moisture, protein, fat, ash, salt, ST, CHO’S, mesophilic, and fungal and yeast counts at three ripening times. For data shown to have a normal distribution, analysis of variance was used applying a Completely Randomized Design (CRD) with an α = 0.05. Comparison of means was performed with the Tukey-Kramer test with an α = 0.05 in the MINITAB 2017 statistical program (Minitab, LLC, Pine Hall Road, PA, USA). On the other hand, when the data did not have a normal distribution, the Kruskal Wallis statistical test was used (17). Comparison of means was performed by the Steel-Dwass-Critchlow-Fligner test with an α = 0.05, data were analyzed in the XLSTAT 2019 statistical software (XLSTAT, LLC, Rue Damrémont, Paris, France).

### Bioinformatics analysis of metagenomics

The quality control of 16S gene sequences and ITS regions was carried out using the MultiFastQC program of the Galaxy platform (18). Adapter removal and subsequent sequence quality checks were performed using the Trimmomatic and MultiFASTQC programs, respectively. Flash v1.2.11 (https://ccb.jhu.edu/software/F LASH/) was used to join paired sequences from amplicons; repeated and chimeric sequences were subsequently removed using the Vsearch program (https://github.c om/torognes/vsearch). Taxonomic profiling was performed using Parallel-Meta Suite version 3.7 (http://bioinfo.single-cell.cn/parallel_meta.html) with the 16S Greengenes database for bacteria and the ITS2 database for fungi and yeasts. Shannon, Simpson and Chao1 indices were calculated using the Phyloseq package of R. Finally, relative abundance (RA) plots and Venn diagrams were generated using Perl and R scripts.

### Physicochemical analysis of Queso Crema de Chiapas

The variables a_w_, protein (%), salt (%), ST (%), CHO’S (%), and aerobic mesophilic counts at the different ripening times presented a normal distribution. On the other hand, the variables moisture, fat, ash, fungal counts, and yeast had a non-normal distribution (data not shown). The results of the statistical analysis are shown in Table 1. Moisture ranged from 33.36 to 34.56 % and fat changed from 35.15 to 36.9 %; these variables had no statistical difference during ripening. This trend is the same as reported in vacuum-packed San Simón da Costa cheese (19), in which the casing prevented water loss, decrease of a_w_ and fat. On the other hand, fat is not modified because the lipolytic activity of LAB is limited because saturated and unsaturated fatty acids have a toxic effect on bacteria (20) and pH 9 is the optimum of lipolytic enzymes (21).

**Table 1.** Physicochemical analysis of Queso Crema de Chiapas at different ripening times.

| Variables | T1 | T2 | T3 |
| --- | --- | --- | --- |
| $a_w$ | 0.93±0.02a | 0.93±0.02a | 0.93±0.01a |
| Moisture (%) | 34.56±0.90a | 33.36±2.48a | 33.98±0.9a |
| Protein (%) | 27.81±0.99a | 25.7±0.38b | 22.87±0.89c |
| Fat (%) | 36.01±0.52a | 35.15±2.43a | 36.9±0.85a |
| Ash (%) | 3.54±9.90a | 3.35±0.75a | 3.58±0.08a |
| Salt (%) | 2.73±0.24a | 1.99±0.11b | 1.91±0.29b |
| Total solids (%) | 69.26±1.90a | 64.61±1.51b | 64.61±1.51b |
| Carbohydrates (%) | 1.91±1.28a | 1.33±0.9a | 1.91±0.29a |

Means ± standard error followed by different letters (a, b, c) in a row indicate statistical difference between times ( $p < 0.05$ ).

In the cheese samples analyzed, a decrease was observed in three variables, which were total solids (TS), protein and salt as shown in Table 1. The TS values decreased from 69.26 to 64.61 %, protein decreased from 27.81 to 22.87%, as did salt (2.73 to 1.91 %). Research on Örgü cheese (22) and Greek Feta cheese (23) showed the same behavior in total solids and protein variables during ripening because degradation reactions decrease protein and total solids content in cheese (22). Additionally, in Feta cheese and in artisanal raw sheep milk cheese, it was found that salt decreased with ripening, and this was attributed to the fact that high pH and fat content prevented the absorption of this compound (24); (25).

### Microbiological analysis of Queso Crema de Chiapas

In the cheeses analyzed across the three times (T1 = 2, T2 = 29 and T3 = 58 d) no growth of total coliforms was observed, which can be attributed to the fact that during ripening antimicrobial compounds, organic acids and ethanol, are generated; this effect is also attributed to the physicochemical changes in the cheese. such as: decrease in pH, a_w_ and moisture (26); (27).

On the other hand, the mesophilic bacteria count had no significant statistical difference in the three ripening times of the Queso Crema de Chiapas, the values ranged from 4.36 to 6.13 log_10_ cfu g^-1^ (Fig 1), which may be associated with the fact that the physicochemical properties were not significantly modified during ripening.

**Fig 1.**
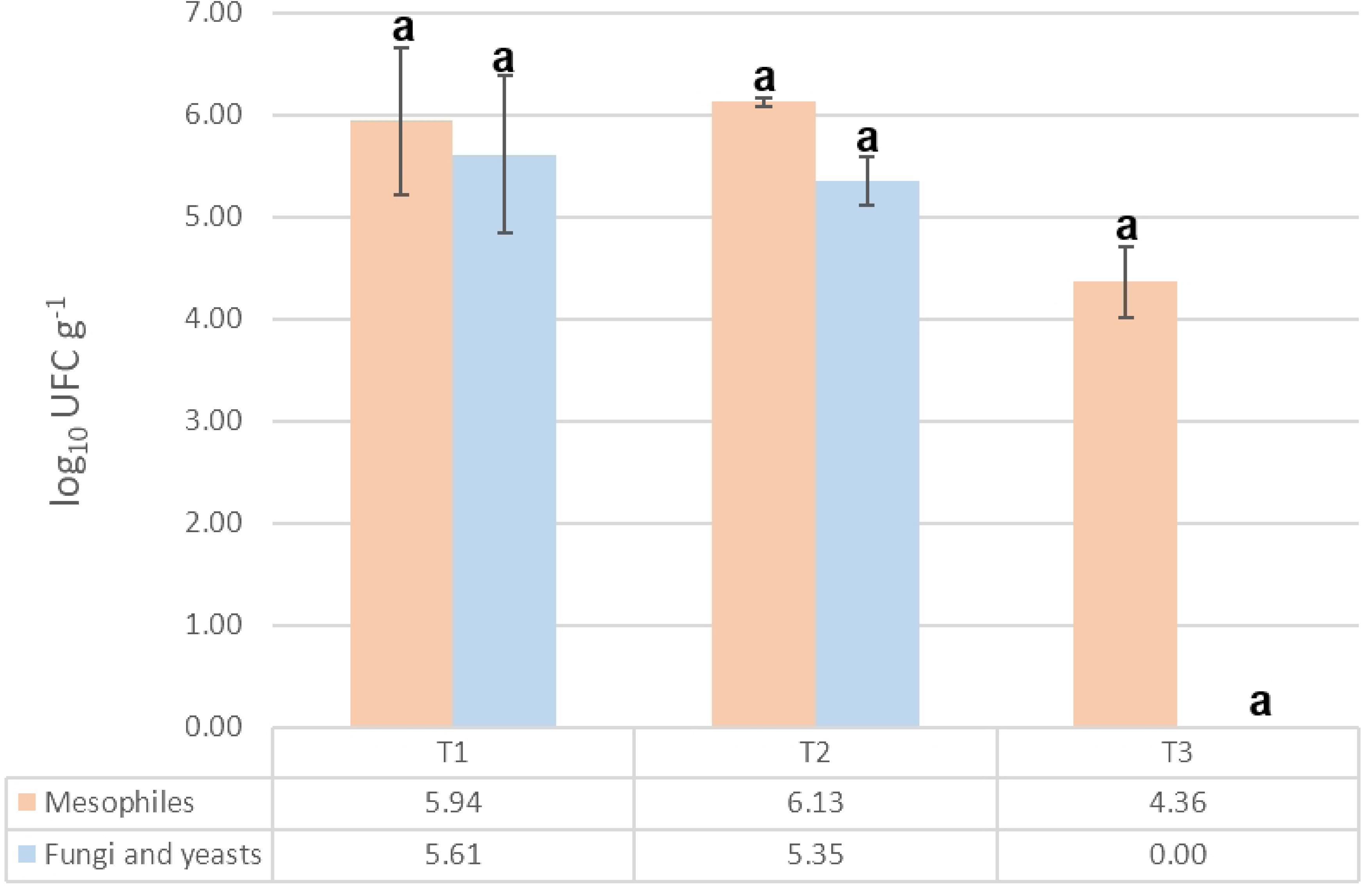
Mesophilic bacteria, fungi, and yeast counts in Queso Crema de Chiapas at three ripening times (T1 = 2, T2 = 29 and T3 = 58 d). Bars with the same letter indicate no significant difference.

Likewise, for fungi and yeasts, no significant difference was observed during the ripening process related to the little variation in the physicochemical properties of the cheese (Fig 1). Some research reports that the presence of fungi in cheese is controversial, because they inhabit the air and some produce toxins (28). In industrial Oaxaca cheese, aflatoxin B1 and aflatoxicol, originating from corn flour, are reportedly added to the cheese dough to thicken it (29). However, the presence of certain genera of fungi and yeasts is known to be important, as they participate in the proteolysis and lipolysis of cheese, contributing to its sensory properties (30).

In this study, the direct relationship between culture media-dependent methods and metagenomics was not determined; however, it was evident that total coliforms were found in low abundance (data not shown) and were not alive. It was also observed that mesophilic microorganisms were present in high quantities and abundances throughout cheese ripening. Additionally, fungi and yeast DNA were observed at T3, however, they were not alive because there was no growth in the culture media.

### Metagenomics

A total of seven genera of bacteria were identified during the cheese ripening time. At T1, *Streptococcus*, *Lactobacillus*, *Aeromonas*, *Lactococcus*, and *Enterobacter* were found. For T2, *Streptococcus*, *Lactobacillus*, *Lactococcus*, *Aeromonas* and *Chryseobacterium* were found. Finally, at T3 a total of six genera were identified: *Stretococcus*, *Lactobacillus*, *Lactococcus*, *Aeromonas*, Bifidobacteri*um* and *Chryseobacterium* (Fig 2). The dynamics in the abundance of the different genera of microorganisms that occur during cheese ripening are related to physicochemical changes and environmental factors, which promote the selection of specific microbial species (31). In the samples analyzed, it was observed that at 58 d the genera increased from five to six; this increase has also been reported in Dutch-type cheese (32).

**Fig 2.**
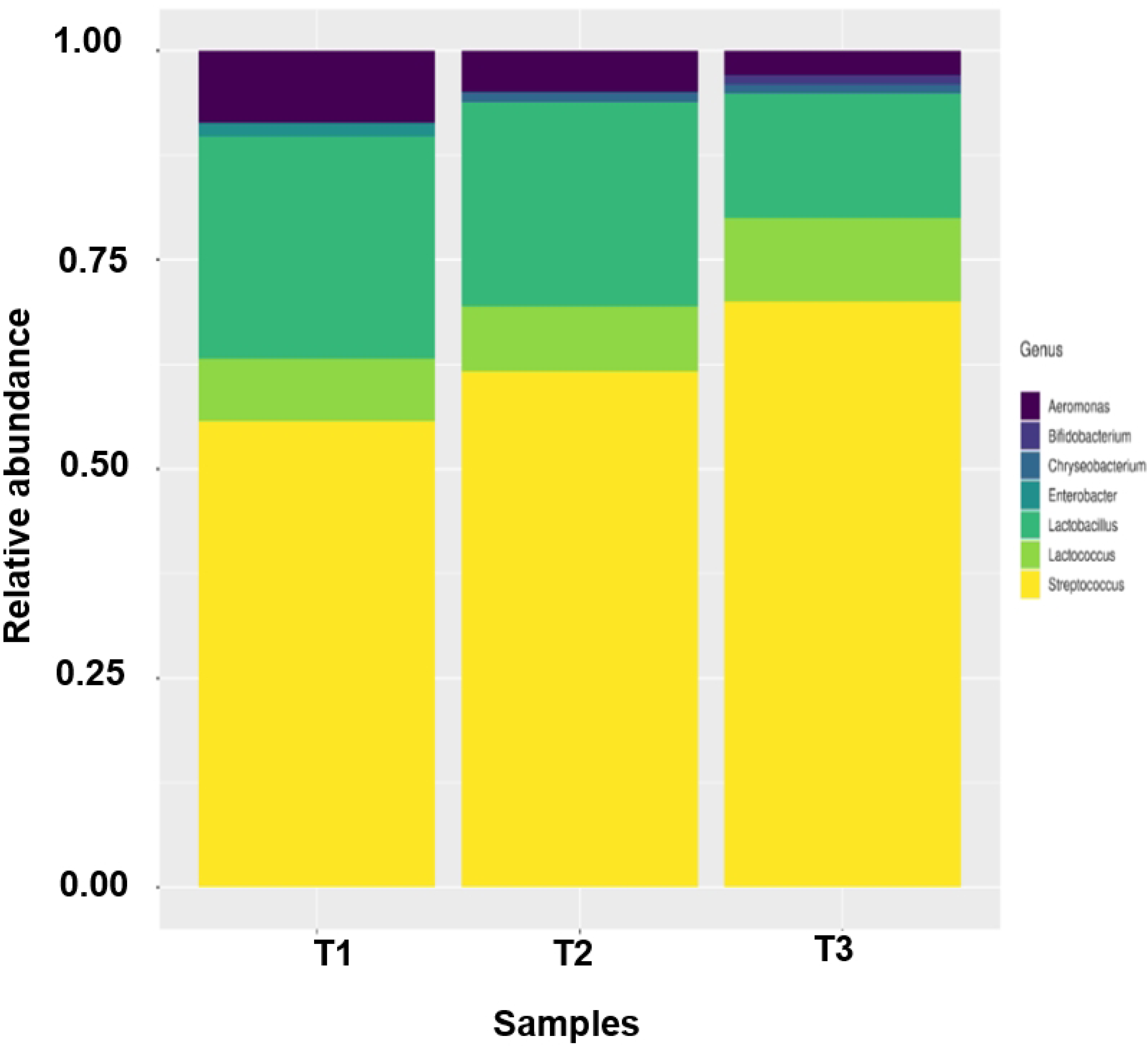
Relative abundance of bacteria in Queso Crema de Chiapas during ripening (T1 = 2 d, T2 = 29 d and T3 = 58 d) considering the V3 and V4 regions of the 16S gene.

Alpha diversity shows the variety of species or richness within a community (33), Shannon’s indices and Simpson’s diversity index are the most commonly used (34). These indicators, along with Chao’s1 are calculated based on species and OUT abundance (35). The cheese with 2 d of processing presented the highest species richness and diversity (Table 2), given that the Shannon value (3.09) has a range between 1.5 to 4.5 (36) and Simpson (0.93) is between 0 and 1, indicating that microbial diversity is high (37). Likewise, it was found that the diversity of the samples decreased along the ripening time; this behavior coincided with that reported in Parmigiano Reggiano cheese, due to the survival of microorganisms that use alternative energy sources to carbohydrates in milk and a low a_w_ (38). With the Chao1 index, an increase in species richness was observed during ripening time in Queso Crema de Chiapas, reaching its highest value of 136.5 at 58 d. This behavior was similar in Canadian Cheddar cheese; in which it is noted that a value higher than 24 in the Chao1 index increases bacterial genera (37).

**Table 2.** Alpha diversity with Chao1, Shannon and Simpson indices at three ripening times (T1 = 2 d, T2 = 29 d and T3 = 58 d) with the 16S gene in Queso Crema de Chiapas.

| Sample | Observations | Chao1 | Shannon | Simpson |
| --- | --- | --- | --- | --- |
| T1 | 43 | 97.167 | 3.092 | 0.927 |
| T2 | 44 | 84.625 | 3.057 | 0.919 |
| T3 | 42 | 136.5 | 2.929 | 0.906 |

The most abundant genus in Queso Crema de Chiapas at the different ripening times was *Streptococcus*, followed by *Lactobacillus* and then *Lactococcus*. This may be due to the presence of low molecular weight nitrogenous compounds and associations with the *Lactobacillus* genus (39). The genera *Streptococcus* and *Lactobacillus* have been reported in Mexican cheeses such as Poro cheese and Bola de Ocosingo cheese (39) because they are naturally found in milk and whey.

In the Queso Crema de Chiapas, the second most abundant genus was *Lactobacillus*, which decreased over time. This behavior coincides with that reported in Tulum cheese (40) and Feta cheese (41) where the amount of salt (7%) and a low pH (4.4) affected the survival of this genus. The change in the abundance of *Lactobacillus* strains with respect to ripening time is due to the modification of physicochemical characteristics such as total solids, protein, and pH, which have been reported in Greek Feta cheese (23) and Parmigiano Reggiano cheese (38).

The genus *Lactococcus* was the third most abundant; at the time of 58 d it had a relative abundance (RA) of 0.25. This behavior coincides with that reported in Gouda cheese (42) and Feta cheese (43) that presented abundances of 58.5% and 90%, respectively. *Lactococcus* is not very strict in terms of temperature, as it grows between 10-40°C and has tolerance to high salt concentrations (44). The genera *Streptococcus*, *Lactobacillus,* and *Lactococcus* can contribute to the sensory properties of Queso Crema de Chiapas throughout the ripening process. In Cantal and Historic Rebel cheese, the genus *Lactobacillus* subsp. was correlated with esters and alcohols such as 2-heptanol, derived from ketone reduction, while *Lactococcus* subsp. was correlated with ketones (45); (46). Esters are associated with fruity, sweet, floral and spicy notes (46) and n-alcohols are also associated with fruity notes which may be the product of fatty acid lipolysis (47); (46).

In the Queso Crema de Chiapas, *Aeromonas*, *Enterobacter*, *Chryseobacterium*, and *Bifidobacterium* were also found in smaller AR. The first genus reaches the food due to poor production practices and decreases during cheese ripening, which is consistent with what was reported in Paipa cheese (48). *Aeromonas* have been identified in Panela cheese, Adobera cheese, Ranchero cheese, Jocoque (49) and Sonora queso fresco (50). Some *Aeromonas* strains have virulence genes that, when combined with virulence genes from other strains, can cause severe diarrhea in humans (51). The *Enterobacter* genus was found at T1 and decreased throughout cheese ripening, this behavior was similar in soft and semi-hard cheeses (52), this mainly due to physicochemical changes during ripening (53).

At times T2 and T3, the genus *Chyseobacterium* was found, which has been reported in dairy products and raw milk by Hugo *et al.* (54); it was also reported in Poro cheese from Tabasco (31). Microorganisms belonging to this genus are emerging opportunistic pathogens, and together with other bacteria are indicators of the production site (55). In milk it has been related to the production of yellow pigments (56) and in Fontina cheese it contributes sensory properties by proteolysis and lipolysis (57).

Another important genus found in the third ripening time was *Bifidobacterium,* which has also been reported in Italian cheese (58), Parmigiano Reggiano cheese (38) and “Tomme d’Orchies” cheese (59). This genus is important due to its association with 11 bioactivities (60). However, it must be found in sufficient quantities, have adherence, and survive intestinal conditions (60).

To visualize the distribution of the 213 OTUs obtained among the different maturation times, a Venn diagram was constructed (Fig 3). The data indicated that 102 OTUs included 48% of the total reads, which were shared among the three maturation times. 36 of the OTUs accounted for 17% and are microorganisms that are shared between two maturation times (T1-T2; T2-T3; T3-T1) and 75 OTUs corresponding to 35% are unique OTUs. The highest number (n=28) of unique OTUs was located at the maturation time of 58 d (T3) and the lowest number (n=22) of OTUs was observed for the time of 2 d (T1). This behavior can be explained because at T3 there is a lower amount of NaCl, which favored the presence of bacteria that can affect feed quality by generating unpalatable compounds (61). Shared OTUs of higher abundance can provide information of possible bacteria that can be selected as starter cultures (62) and also, of pathogenic bacteria that prevail throughout ripening. However, the more time-specific bacteria contribute to unique characteristics of the cheese at each ripening time.

**Fig 3.**
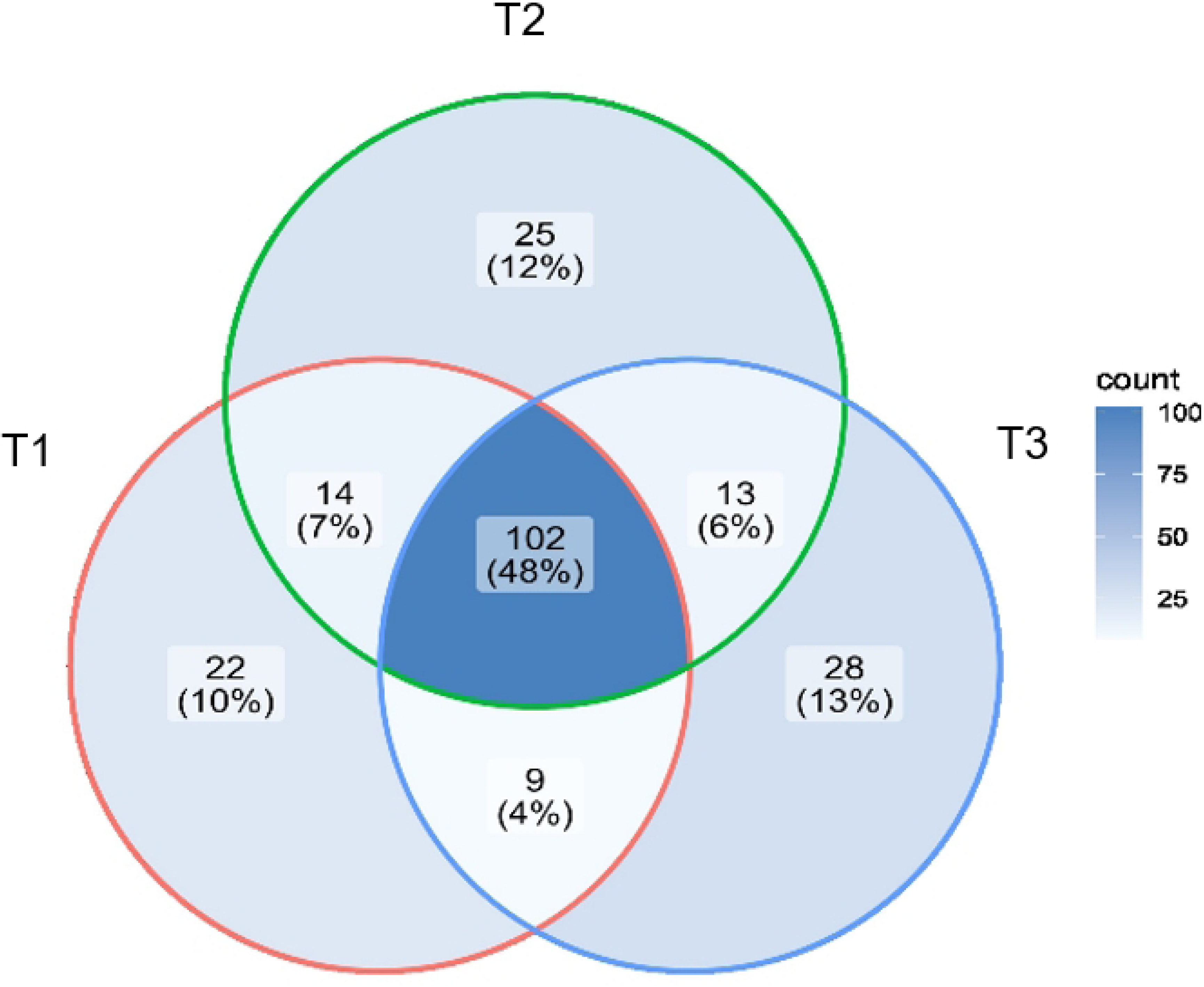
Venn diagram showing the number of specific and common OTU between three ripening times of Queso Crema de Chiapas.

### Fungal and yeast dynamics in Queso Crema de Chiapas

Table 3 shows that the Shannon, Simpson and Chao1 indices showed that as ripening time increases, the diversity and richness of fungal and yeast species increases, reaching their maximum value: Shannon = 3.394, Simpson = 0.955 and Chao1 = 144.75 at 58 d. In the study by Dimov *et al.* (63), they reported that, in Bulgarian green cheese, the values of the Shannon and Chao1 indices also increased with ripening time. Levante *et al.* (64) indicated that microbial diversity in buffalo Mozzarella cheese was modified according to the place of production.

**Table 3.**
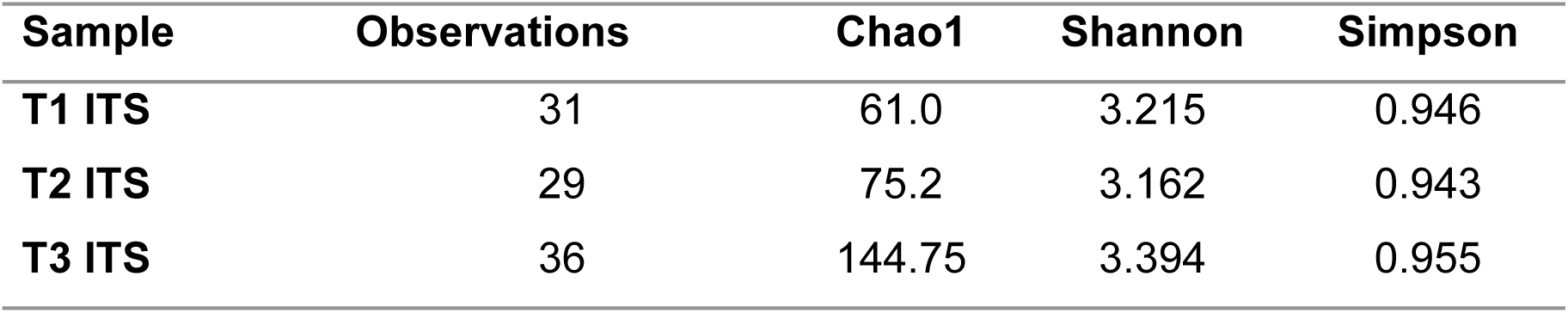
Alpha diversity with Chao1, Shannon and Simpson indices at three maturation times (T1 = 2, T2 = 29 and T3 = 58 d) determined with ITS region sequencing.

The most common genera found in different cheeses are *Candida*, *Pichia*, *Saccaromyces*, and *Trichosporon* (65). Sixteen fungal genera have been identified in Mexican cheeses and the most abundant were *Galactomyces, Saccharomyces*, and *Scheffersomyces* (49).

In Queso Crema de Chiapas, eight species of fungi and two yeasts were identified throughout the ripening process. In T1, the genera *Saccharomyces cerevisiae* (0.47 AR), *Candida versatilis* (0.5 AR), *Candida tropicalis* (0.016 AR) and *Candida etchellsii* (0.014 AR) were found. In T2 the most abundant species was *C. versatilis* (0.88 AR); in addition, *Candida kruissii* (0.062 AR), *C. etchellsii* (0.03 AR), *Phaeoacremonium hungaricum* (0.013 AR) and unclassified *Pichia* (0.025 AR). In T3 the most abundant species was *C. versatilis* (0.625 AR); other species identified were *C. kruisii* (0.0625 AR), *C. etchellsii* (0.125 AR), *C. tropicalis* (0.042 AR), *P. hungaricum* (0.042 AR), *Pichia* (unclassified) (0.042 AR), *Trichosporon debenurmannianum* (0.022 AR), *Malassezia furfur* (0.022 AR), *S. cerevisiae* (0.011 AR) and *Candida parapsilosis* (0.011 AR).

A greater number of fungal species were observed as the ripening time increased, this behavior is similar to that reported in Cantal cheese (45) and Austrian Vorarlberger Bergkäse cheese (35), the difference in species was due to the place from which the cream that was added was obtained, the presence of polyunsaturated fatty acids and the presence of n-alcohols and the facilities (35); (45).

At the three ripening times of Queso Crema de Chiapas, the three most abundant strains of fungi were *C. versatilis*, *C. etchellsi*, and *C. tropicalis*. In the cheese analyzed, the *C. versatilis* strain, in addition to being the most abundant, was the most variable during ripening. In Vorarlberger Bergkäse cheese, the genus *Candida* was abundant at the beginning of ripening and decreased over time (35). The *C. versatilis* strain has been identified in whey, yogurt, and cheeses (66), such as Gorgonzola and Danish (67); *C. versatilis* is a salt-resistant strain; in cheese it is unknown how it acts on certain substrates; however, using glucose it produces glycerol and mannitol (68). *C. etchelsi* is the second most abundant strain and whose abundance decreased as ripening progressed. This strain has been reported in Cotija cheese (69) and in Taleggio cheese with a ripening period of seven days (70). The contribution of *C. etchelsi* is not fully explained in cheese, however, it was found that in a Chinese bean chili paste they are able to contribute to the synthesis of esters that contribute to flavor (71).

*C. tropicalis* is the third most abundant strain and increased with ripening time. This strain was identified in Oaxaca cheese (72), raw bovine milk, in Cremoso cheese (73) and in cheese Mihalic (74). Dinika *et al.* (75) reported that in the fermentation of Mozzarella cheese whey with *C. tropicalis*, peptides were hydrolyzed, generating amino acids such as Asp, Glu, Thr, Val, Ile, and Lys, related to the biosynthesis of glutamate as an amine donor.

*S. cerevisiae* presented the highest AR value (0.47) in Queso Crema de Chiapas, in T1 (Figure 4). This yeast has been identified in Frescal cheese (76), in Gorgonzola cheese and in Danbo cheese as a producer of aldehydes and branched alcohols (77). *Saccharomyces* has also been reported by Murugesan *et al.* (49) in Mexican cheeses such as: Adobera, Doble Crema de Chiapas, Ranchero, Chihuahua, Cincho, Oaxaca, Canasta, Manchego and Jocoque.

**Fig 4.**
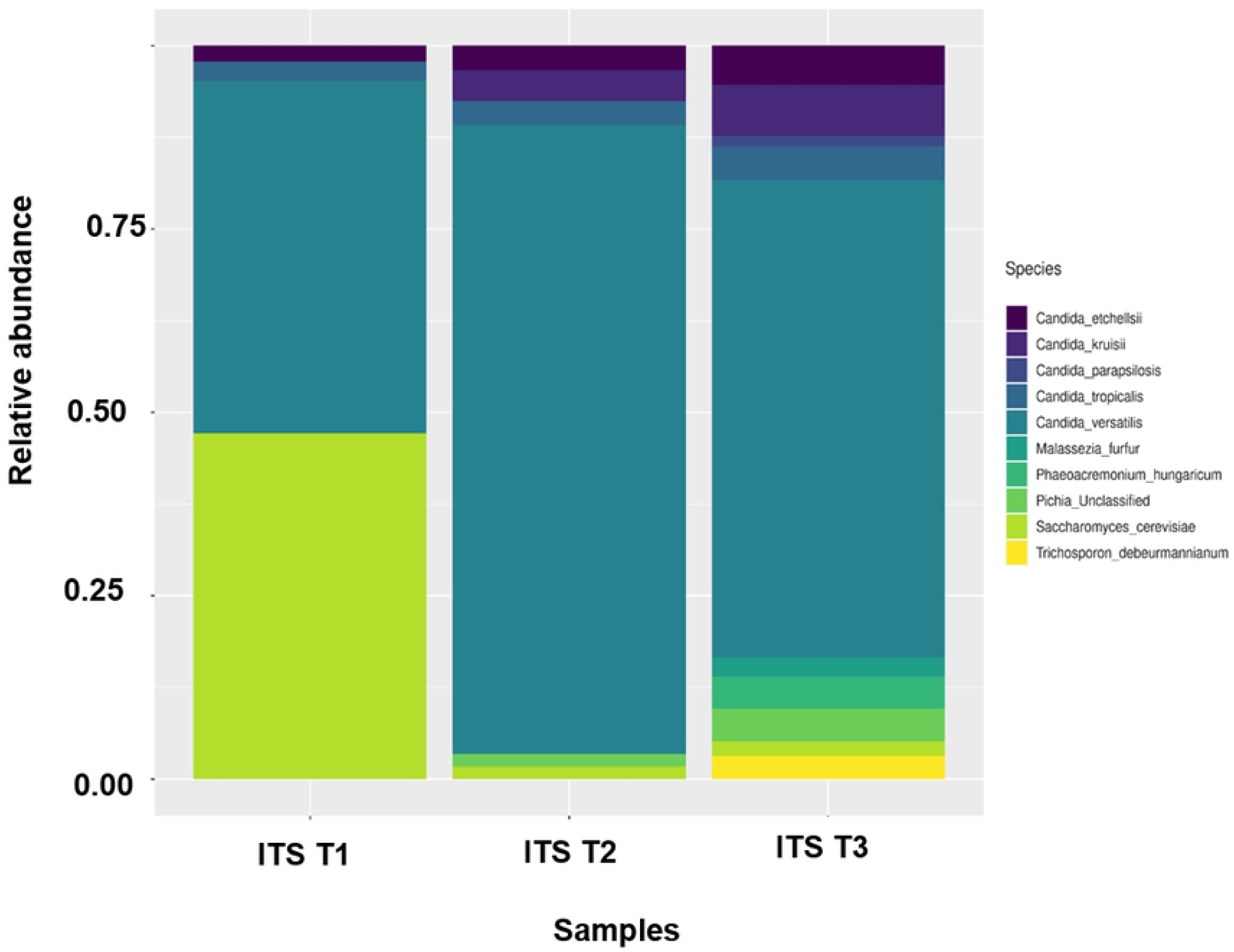
Relative abundance of fungi and yeasts of Queso Crema de Chiapas during ripening (T1=2, T2=29 and T3=58 d) detected by sequencing the ITS1 and ITS2 regions.

Other strains were found in lower abundance in the Queso Crema de Chiapas samples such as: *C. kruisii, P. hungaricum, Pichia* (unclassified), *T. beermannianum, M. furfur*, and *C. parapsilosis*. The *C. kruisii* species decreases as maturation time passes. *P. hungaricum* was found in T3 of ripening in Queso Crema de Chiapas; this genus has not been reported in cheeses. However, *P. hungaricum* has been isolated from the environment, diseased woody plants, wood, grapevines, humans with pheohyphomycotic infections, bark beetle larvae, arthropods, and soil (78).

The *Pichia* genus was found in the third maturation time with an AR of 0.06 (Fig 4). This behavior was opposite to what was reported in Fossa cheese, in which *Pichia occidentalis* was found in the environment where the cheese was made and decreased with ripening (79). The genus *Pichia* has been identified in Mexican artisanal cheeses (49), in Cotija cheese (69) and in Roquefort cheese (80). The genus *Pichia* has been used as a starter culture in Cantal cheese (81); in Kasajo cheese it was observed that depending on the strains different aroma profiles are obtained, for example, the presence of *Pichia kudriavzevii* A11 was related to the presence of alcohols, acetic acid, and acetates (82).

In Queso Crema de Chiapas, *T. debeurmannianum* was found at maturity time T3. This genus was reported in raw milk and Quebec cheese (83). Its presence indicated contamination, because *T. debeurmannianum* has been identified in clinical samples from patients with urinary tract infections (84), diabetic foot infection (85), fungal ampoules (86); as well as in the environment, skin, intestinal tract, and vagina (87). The genus *Trichosporon* utilizes different carbohydrates, carbon sources and degrades urea, but is not considered a fermentative microorganism (87).

In the cheese studied, *M. furfur* was found at T3 (Fig 4), which is not common in cheese; however, the genus *Malassezia* is lipid-dependent (88). Because this genus grows in aerobic or anaerobic conditions from 33 to 41 °C (89) it is an inhabitant of human skin (90) and breast milk (91); in dairy products it has not been investigated enough to be an indicator of contamination.

*C. parapsilosis* in the cheese analyzed was identified in low abundance (0.031) at T3. The *C. parapsilosis* strain favored the growth of *Lactobacillus paracasei* in Comté cheese (92) and has been identified in Cheddar cheese, Swiss-type cheese, and American blue cheese (93). Their presence favors the hydrolysis of α-lactoalbumin and β-lactoglobulin and they produce mainly alcohols (92).

To evaluate the distribution of the 72 fungal and yeast OTUs obtained among the different maturation times, a Venn diagram was constructed (Fig 5). The data indicated that 30 OTUs included 42% of the total reads, which were shared across the three maturation times. 10 OTUs are common in two maturation times and account for 14% of the total reads. 32 OTUs which are 44% of the total reads are spread over the three times individually. The highest number (n=13) of unique OTUs was located at maturation times T1 and T3, respectively, and the lowest number (n=6) of unique OTUs was observed for T2. The number of fungal species at times T1 and T3 only differed by one species, this can be explained to some fungi having hosmotolerance; for example, *S. cerevisiae* has been reported to produce glycerol to tolerate stress conditions (94). On the other hand, it was shown that the number of species is lower at T2; since there were no modifications in the physicochemical characteristics, we could suppose that the bacteria present at this time could have inhibited certain fungal strains. Leyva-Salas *et al.* (95) found that organic acids produced by *Lactobacillus* strains inoculated in semi-hard cheese and yogurt presented antifungal activity against *Penicillium*, *Candida* and *Trichosporon*.

**Fig 5.**
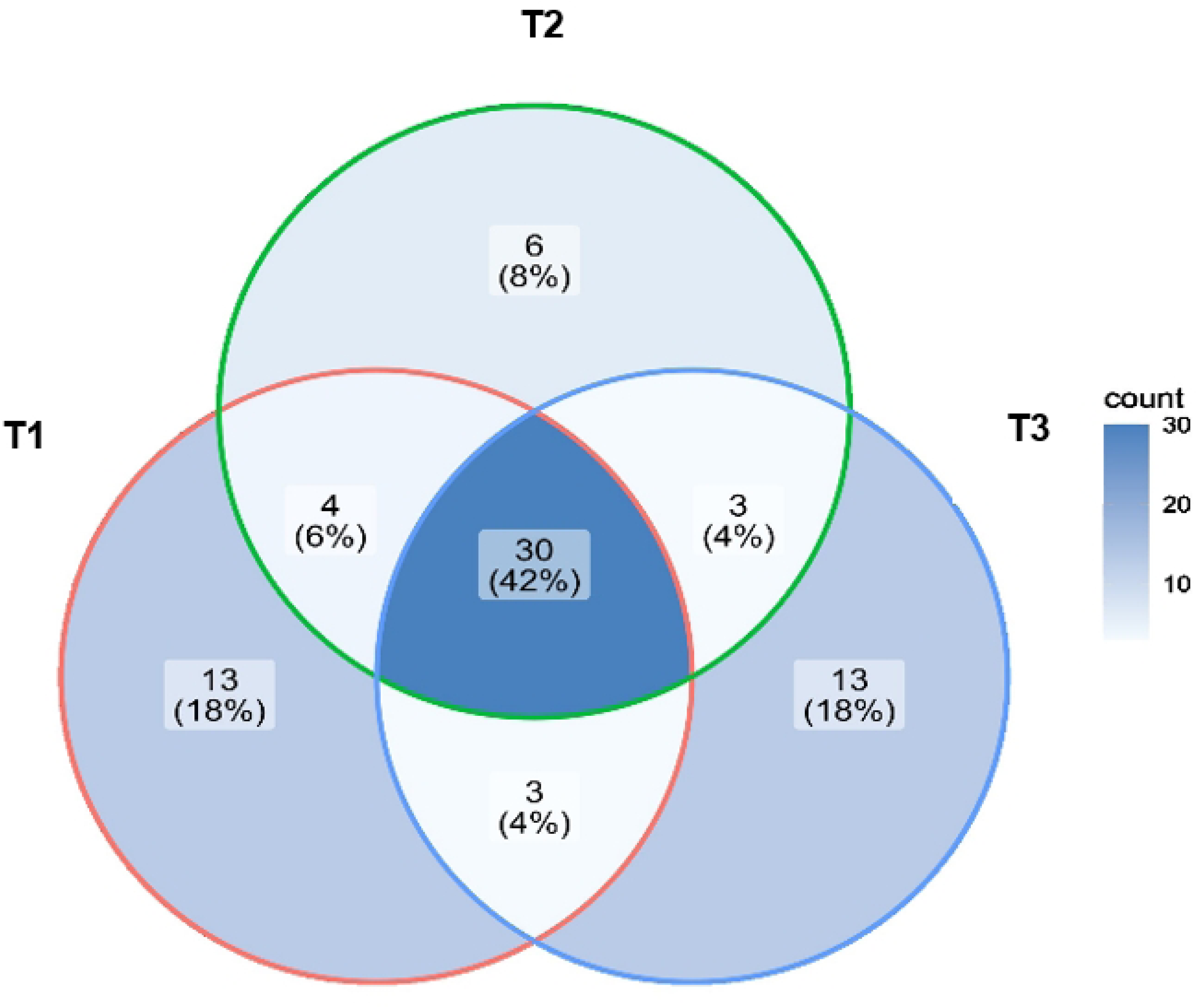
Venn diagram showing the number of specific and common OTUs (OTU 30) between three maturation times of Queso Crema de Chiapas raw cow’s milk (2, 29 and 58 d).

## Conclusions

Our study reveals the microbial dynamics of the artisanal Mexican cheese “Queso crema de Chiapas” based on a metagenomic analysis. Three distinct maturation stages (2, 29, and 58 days) were examined, and changes in both microbial and fungal communities were observed, with the most abundant organisms being *Streptococcus*, *Lactobacillus*, *Lactococcus*, *Candida versatilis*, *Candida etchellsi*, and *Candida tropicalis*. This allowed us to understand the impact of their interactions as well as their influence on texture and flavor at each stage. We identified specific species, such as the aforementioned *Candida strains*, which distinguishes our work from studies that focus on broader taxonomic groups. Although some species common to the production of other cheeses worldwide were detected, the use of raw cow’s milk, the extended fermentation process, and the unique conditions in Mexico resulted in a microbial and fungal community exclusive to the cheese under study. Additionally, our findings revealed the presence of potentially pathogenic organisms, such as *Aeromonas* and *Chryseobacterium*, setting the stage for future research to explore growth conditions and prevalence and to implement targeted control measures to further reduce or eliminate them. Our work represents a significant advance in understanding the maturation process of artisanal cheeses and underscores the importance of integrating metagenomic techniques with traditional culture methods.

## Acknowledgments

We thank the Consejo Nacional de Humanidades Ciencias y Tecnologías (CONAHCYT) for the grant 2020-000013-01NACF-03858 awarded to Blanca Nayelli Ocampo Morales. We also thank Jerome Verleyen for his technical support and for granting us access to the HPC infrastructure at the Unidad Universitaria de Secuenciación Masiva y Bioinformática, Instituto de Biotecnología (UNAM), and Mabel Rodríguez for her invaluable assistance with proofreading, and insightful suggestions.

## Authoŕs contribution

Blanca Nayelli Ocampo Morales: Formal analysis, Methodology, Original draft. Ernestina Valadez Moctezuma: Conceptualization, Formal analysis, Methodology, Drafting-revision. Karel Johan Estrada Guerra: Formal analysis, supervision, validation, Software, Drafting-revision. Arturo Hernández Montes: Data collection, Project management, Proofreading and editing.

## Financing information

Dirección General de Investigación y Posgrado (DGIP; State of Mexico, Mexico) research projects 22004-DTT-62 and 23001-DTT-62.

## Data availability statement

Data supporting the conclusions of this study are available from the corresponding author upon request.

